# Getting to the bottom of social learning: Chimpanzees copy arbitrary behavior from conspecifics

**DOI:** 10.1101/2024.08.30.607436

**Authors:** Edwin J. C. van Leeuwen, Emile Bryon, Alex Rogers, Aurore Balaran, Peggy Motsch, Jake S. Brooker

## Abstract

Studying animal culture has been crucial for understanding the complexities of knowledge transmission and tracing human culture’s evolutionary origins^1^. Defined as the use of tools to provide clear practical benefits to individuals, well-documented examples of material culture include nut-cracking^2^ and termite fishing^3^ in chimpanzees. Additionally, there is growing interest in animal social traditions, which appear crucial for social interaction and group cohesion. We have previously documented such a tradition, in which chimpanzees copied inserting blades of grass in their ears from one persistent inventor^4^. Now, over a decade later, we have observed an unrelated group of chimpanzees where 5/8 individuals began wearing grass in their ears and 6/8 from their rectums. As of 2024, one newly introduced chimpanzee has adopted the grass-in-ear behavior. Given that the behaviors were not observed in seven other groups in the same sanctuary (*N*=148), we conclude that social learning of arbitrary behavior occurred and discuss our findings considering the larger scope of animal culture.

## Main text

Documenting seemingly non-functional behaviors is essential as they offer parallels to aspects of human culture, where many social customs and traditions serve more to reinforce group identity and cohesion than to impart specific skills. Behaviors such as fashion trends, rituals, and social norms in human societies often do not have direct material benefits but are critical for social bonding and cultural identity. Similarly, vocal traditions in humpback whales^5^, complex social games in capuchin monkeys^6^, and handclasp grooming in the *Pan* apes^7^ indicate that social traditions foster strong affiliative bonds and group cohesion, highlighting the possible importance of non-material culture in facilitating group identity and social integration in animals.

Now, we report the emergence and diffusion of two non-instrumental behaviors in a group of semi-wild chimpanzees at Chimfunshi Wildlife Orphanage Trust (Zambia). First, in 2023, chimpanzees started to engage in the grass-in-ear behavior (GIEB), which we defined as *“*inserting grass or sticks in one’s ear and leaving it hanging without manual assistance” (Figure 1a). On August 16^th^, the first chimpanzee (Juma) engaged in GIEB, followed by four more individuals within one week (Figure 1a). Upon the integration of three immigrants into this group in December 2023, one (Aimée-Love) adopted GIEB by April 2024.

**Figure 1.**
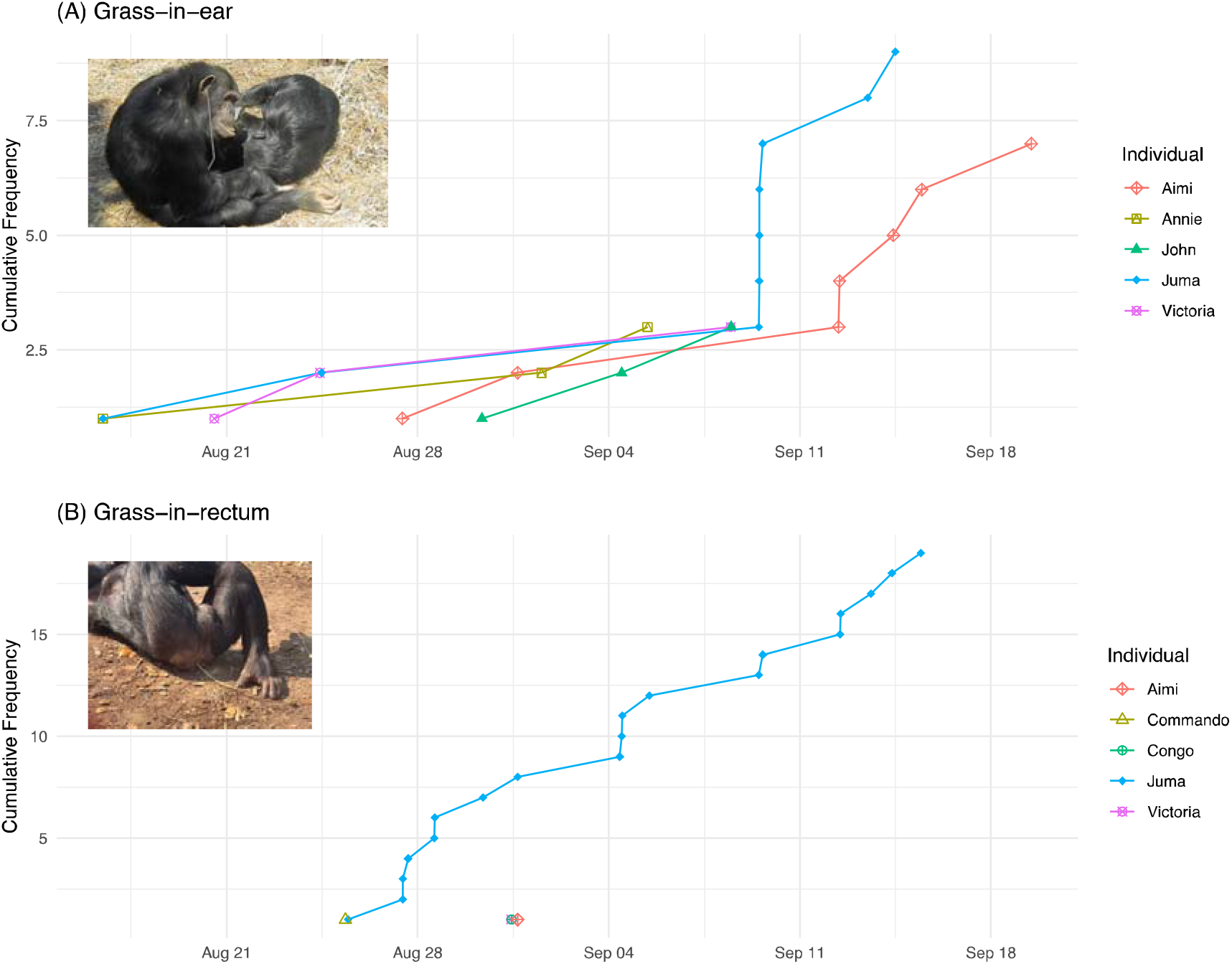
Cumulative frequencies of (a) grass-in-ear adopters, and (b) grass-in-rectum adopters in one group of chimpanzees at Chimfunshi (“Sudan” group) in 2023.

Second, on August 27^th^ (2023), Juma engaged in a GIEB variation, namely the grass-in-rectum behavior (GIRB), which we defined as “putting grass or sticks in one’s rectum and leaving it hanging without manual assistance” (Figure 1b). On the same day, we observed Commando engaging in GIRB. Aimi, Congo, and Victoria subsequently engaged in GIRB (all on 31 August 2023; Figure 1b). Later in the season (October 7^th^), John engaged in GIRB. Importantly, in the other seven groups comprising *N*=148 chimpanzees, during approximately 630 observation hours between May 2023 and August 2024, we only observed two other chimpanzees engage in GIEB (in Group 4), and none in GIRB.

To investigate whether these behaviors were socially transmitted, we applied Network-Based Diffusion Analysis (NBDA)^8^. We constructed a social network based on group membership, with ‘1’ indicating chimpanzees belonging to the same group and ‘0’ otherwise. Sex and age were included as individual-level variables (ILVs) that potentially influence the acquisition order. We fitted models incorporating all combinations of ILV effects and employed multi-model inference using Akaike’s Information Criterion corrected for sample size (AICc) to assess the support (total Akaike weight) for social versus asocial transmission through the network.

For both the diffusion of GIEB and GIRB, the group network received the most support (both 99%), with asocial models supported for less than 1% (corrected for the number of models fitted). Where *s* is the rate of social transmission per unit connection relative to the asocial rate for females, GIEB yielded 95%CI of *s*=0.99-7.00 and GIRB 95%CI of *s*=0.99-8.99, corresponding to an estimated 83.6% and 100% of chimpanzees having learned the behavior via social learning, respectively. Age and sex did not obviously impact the rates of asocial or social learning (all estimates < 12% likely to be in the best model).

We document the second independent wave of an arbitrary non-functional behavior in chimpanzees which spread within three weeks to most of the group after the initial invention (Figure 1a). After a GIEB resurgence (cf. ^4^), a new variant emerged where chimpanzees began wearing grass from their rectums (Figure 1b), also spreading to most of the group within three weeks. Though we primarily observed GIRB by the same individual, the behavior did spread to five other individuals. None of the 148 chimpanzees across the other seven groups performed these grass-related behaviors between May 2023 and August 2024, besides two males in the group where the first wave of GIEB was observed^4^. Moreover, tailored Network-Based Diffusion Analysis provides strong evidence for social over asocial learning. Our findings show that chimpanzees socially learn non-instrumental behaviors from one another, which is scarcely documented in animals.

The precise form of social learning of these behaviors is impossible to identify. The “Zone of Latent Solutions” (ZLS) hypothesis is concerned with the difference in social learning capacity between humans and other animals, and posits that animals – unlike humans – do not copy each other, but instead learn individually within an environment that allows for behaviors to manifest^9^. Thus, the ZLS would advance that GIEB and GIRB are latent solutions in chimpanzees and that they could be triggered by social demonstrations, in the absence of the apes copying from others. While it might be true that chimpanzees could invent the GIEB and GIRB by themselves, and thus that theoretically they do not *need* to copy the behavior, the fact that they do it so quickly (for the first time) in response to others doing it points to them copying the behavior anyway. When humans become smokers, do they not copy the smoking-related-behaviors from others, even though all relevant actions could easily be invented by everyone? Here, we also note that both GIEB traditions could have emerged from the originators *copying* caregivers cleaning their ears with matchsticks (*pers*.*comm*. Dominique Chinyama). Hence, we question whether the ZLS hypothesis may be too anal with respect to dismissing copying in other animals and propose to cast the “copying net” wider, especially for spontaneously emerging behaviors.

Non-instrumental behaviors like GIEB and GIRB do not necessarily represent “solutions” to problems, which the Zone of Latent Solutions account implies is a prerequisite for discussing culture. Such behaviors may reveal chimpanzees’ latent capacity to copy behaviors from one another, akin to human trends. The evolution of cultural copying in humans may have been influenced by factors beyond instrumental utility, such as homophily and group belongingness^10^. The desire to conform and strengthen social bonds likely played a crucial role in the propagation of behaviors within groups. This suggests that the evolution of our cultural copying capacity might have been driven as much by the social domain as by the need to transmit instrumental skills, underlining the complex interplay between social dynamics and cultural evolution.

## Acknowledgements

We thank the staff of Chimfunshi Wildlife Orphanage Trust, particularly caregivers of the Sudan group including Dominique Chinyama, Reagan Muyanga, Webby Pupe, and the late Emmanuel Chomba. Special thanks also to Jasmin Keizer, Tarini D’Souza, Casey O’Brien, María Laura Cordonet Castagneto, and Musonda Chibwe for their observations. Additional thanks to Zoë Goldsborough for helpful comments on this manuscript. We conducted observations during fieldwork for a project funded by the Templeton World Charity Foundation. E.J.C.v.L. was additionally funded by the European Union under European Research Council Starting Grant no. 101042961—CULT_ORIGINS. The views and opinions expressed are those of the authors only and do not necessarily reflect those of the European Union or the European Research Council Executive Agency. Neither the European Union nor the granting authority can be held responsible for them. The funders had no role in study design, data collection and analysis, decision to publish or preparation of the manuscript.

## Data availability

All videos and data are accessible via an OSF folder at the following link: https://osf.io/wst56/

## Ethics

Data collection comprised purely naturalistic observations and adhered to the legal requirements of Zambia, the International Primatological Society’s Principles for the Ethical Treatment of Nonhuman Primates, and the Chimfunshi Research Advisory Board.

## References

1. Whiten, A. (2021). The burgeoning reach of animal culture. Science 372, eabe6514. 10.1126/science.abe6514.

2. Luncz, L.V., Mundry, R., and Boesch, C. (2012). Evidence for Cultural Differences between Neighboring Chimpanzee Communities. Current Biology 22, 922–926. 10.1016/J.Cub.2012.03.031.

3. Sanz, C., Morgan, D., and Gulick, S. (2004). New insights into chimpanzees, tools, and termites from the Congo Basin. Am Nat 164, 567–581. 10.1086/424803.

4. van Leeuwen, E.J.C., Cronin, K.A., and Haun, D.B.M. (2014). A group-specific arbitrary tradition in chimpanzees (Pan troglodytes). Animal Cognition 17, 1421–1425.

5. Zandberg, L., Lachlan, R.F., Lamoni, L., and Garland, E.C. (2021). Global cultural evolutionary model of humpback whale song. Philosophical Transactions of the Royal Society B 376. 10.1098/RSTB.2020.0242.

6. Perry, S., Baker, M., Fedigan, L., Gros-Louis, J., Jack, K., MacKinnon, K.C., Manson, J.H., Panger, M., Pyle, K., Rose, L., et al. (2003). Social conventions in wild white-faced capuchin monkeys - Evidence for traditions in a neotropical primate. Current Anthropology 44, 241–268. 10.1086/345825.

7. McGrew, W.C., Marchant, L.F., Scott, S.E., and Tutin, C.E.G. (2001). Intergroup differences in a social custom of wild chimpanzees: The grooming hand-clasp of the Mahale Mountains. Current Anthropology 42, 148–153.

8. Hoppitt, W. (2017). The conceptual foundations of network-based diffusion analysis: choosing networks and interpreting results. Philosophical Transactions of the Royal Society B: Biological Sciences 372, 20160418. 10.1098/rstb.2016.0418.

9. Tennie, C., Bandini, E., van Schaik, C.P., and Hopper, L.M. (2020). The zone of latent solutions and its relevance to understanding ape cultures. Biology & Philosophy 35, 55. 10.1007/s10539-020-09769-9.

10. Haun, D., and Over, H. (2015). Like me: A homophily-based account of human culture. In Epistemological Dimensions of Evolutionary Psychology, T. Breyer, ed. (Springer New York), pp. 117–130. 10.1007/978-1-4939-1387-9_6.

